# Transcriptomic Organization of Human Posttraumatic Stress Disorder

**DOI:** 10.1101/2020.01.27.921403

**Authors:** Matthew J. Girgenti, Jiawei Wang, Dingjue Ji, Dianne Cruz, Traumatic Stress Brain Research Study Group, the Million Veteran Program, Murray B. Stein, Joel Gelernter, Keith Young, Bertrand R. Huber, Douglas E. Williamson, Matthew J. Friedman, John H. Krystal, Hongyu Zhao, Ronald S. Duman

**Affiliations:** Department of Psychiatry, Yale University School of Medicine, 34 Park Street, New Haven, CT 06520, USA; Psychiatry Service, VA Connecticut Health Care System, West Haven, CT, 06516; National Center for PTSD, U.S. Department of Veterans Affairs; Department of Biostatistics, Yale School of Public Health, New Haven, CT, 06510, USA; Program of Computational Biology and Bioinformatics, Yale University, New Haven, CT, 06510, USA; Department of Psychiatry and Behavioral Sciences, Duke University Medical Center, 300 North Duke St, Durham, North Carolina, 27701 USA; VA San Diego Healthcare System, La Jolla, CA, 92161; Departments of Psychiatry and of Family Medicine and Public Health, University of California, San Diego, La Jolla, CA 92093; Baylor Scott & White Psychiatry, Temple, TX, USA; Department of Psychiatry, Texas A&M College of Medicine, College Station, TX, USA; Department of Veterans Affairs, VISN 17 Center of Excellence for Research on Returning War Veterans, Waco, TX, 76711, USA; Central Texas Veterans Health Care System, Temple, TX, 76504, USA; VA Boston Healthcare System, Boston, MA, USA; Boston University Alzheimer’s Disease Center and CTE Center, Department of Neurology, Boston University School of Medicine, Boston, MA, USA; Durham VA Healthcare System, 508 Fulton St, Durham, North Carolina, 27705 USA; Department of Psychiatry, Geisel School of Medicine at Dartmouth, Dartmouth Hanover, NH, USA

**Keywords:** Posttraumatic Stress Disorder, Major Depressive Disorder, Transcriptomics, Genomics, Prefrontal Cortex

## Abstract

Posttraumatic stress disorder (PTSD) affects approximately 8% of the general population, with higher rates in extreme stress groups, including combat veterans or victims of sexual assault. Despite extensive study of the neurobiological correlates of PTSD, little is known about its molecular substrates. Here differential gene expression and network analyses of 4 prefrontal cortex (PFC) postmortem subregions of male and female PTSD subjects demonstrates extensive remodeling of the transcriptomic landscape. The data revealed a highly connected down-regulated set of interneuron transcripts in the most significant gene network associated with PTSD and integration of this data with genotype data from the largest PTSD GWAS identified the interneuron synaptic gene *ELFN1* as conferring significant genetic liability for PTSD. We also identified marked sexual dimorphism in the transcriptomic signatures that could contribute to the higher rates of PTSD in women. Comparison with a matched major depressive disorder (MDD) cohort revealed significant divergence between the molecular profiles of subjects with PTSD and depression despite their high comorbidity. Our analysis provides convergent systems-level evidence of genomic networks within the PFC that contribute to the pathophysiology of PTSD in humans.

## INTRODUCTION

Posttraumatic stress disorder (PTSD) is a persistent, maladaptive response to extreme stress that is characterized by re-experiencing, avoidance, and hyperarousal symptoms that can endure for life ^1, 2^. PTSD has a prevalence of approximately 8% in the general population, but rates increase to 25-35% within groups who have experienced severe psychological stress, such as combat veterans, refugees, and assault victims ^3–5^. The risk factors for developing PTSD are multifactorial ^6–12^, and include history of prior stress exposure, level of social support, and premorbid psychological traits. PTSD is 30-40% heritable ^13–18^ and recent studies provide evidence of genome-wide significant risk loci associated with PTSD ^18–20^.

Epidemiological studies report that women are twice as likely to develop PTSD, as well as other anxiety-related disorders compared to men ^21, 22^. Women also experience more severe, debilitating, and persistent symptoms, suggesting the formation of sex-specific neural alterations ^23–25^. Similarly, there are sex differences in fear learning and extinction in both humans and rodents ^26, 27^, and activation of the brain circuitry underlying the fear response, including the prefrontal cortex (PFC), amygdala, and hippocampus is sexually dimorphic, potentially contributing to higher sensitivity in females ^28–30^.

Studies of the molecular underpinnings of PTSD have been limited largely to white blood cells, identifying altered expression levels of pro-inflammatory cytokines and genes related to glucocorticoid activity^31–33^. Here we report the first transcriptome-wide analysis of gene expression changes in postmortem brain of a large cohort of PTSD subjects. The results demonstrate extensive transcriptional changes in 4 discrete PFC subregions, with both combined sex as well as sex specific PTSD transcriptome profiles. Consistent with recent genetic studies of PTSD, the global transcriptomic signatures for PTSD more closely resemble schizophrenia, autism, and bipolar disorder, indicative of some degree of convergent molecular pathologies. Transcriptome-wide association studies using the Million Veteran PTSD GWAS dataset identified pathogenic genes that are associated with PTSD risk and illness state. The results also show that despite the high rates of comorbidity, the transcriptomic signatures of PTSD and MDD differ significantly. These findings provide the first complete genomic analysis of human PFC in PTSD, demonstrating that psychological trauma profoundly alters the molecular landscape of the brain.

## RESULTS

### Transcriptomic profiling of PFC in PTSD subjects

RNA sequencing (RNA-seq) was used to characterize the transcriptomes of four PFC subregions from postmortem tissue of subjects diagnosed with PTSD and matched healthy controls (**Table S1**): dorsolateral PFC (dlPFC; BA9/46), medial orbitofrontal cortex (OFC; BA11), dorsal anterior cingulate (dACC; BA24), and subgenual prefrontal cortex (sgPFC; BA25). These regions were chosen based on functional evidence, including reports of decreased volume in PTSD subjects ^34^ and have been implicated in emotional dysregulation by functional neuroimaging studies ^35^. In addition, dlPFC has been targeted using transcranial magnetic stimulation as a treatment in both disorders^36^. For each brain region we examined 52 PTSD subjects (26 males, 26 females), 45 MDD (18 females, 27 males), and 46 control (20 females, 26 males). Comparison of the combined male and female control and PTSD cohorts identified 393 differentially expressed genes (DEGs) in the dlPFC, 170 DEGs in the OFC, 74 DEGs in the dACC, and 1 in the sgPFC (FDR <0.05) (**Fig. 1A**). Additionally, significant differential regulation in PTSD of individual exons was observed from a total of 1356 genes in the dlPFC, 6862 genes in the OFC, 72 genes in the dACC, and 37 genes in the sgPFC (FDR<0.05). There was also significant differential exon-exon junction usage from 12 genes in the dlPFC, 3336 genes in the OFC, 1 gene in the dACC, and 0 genes in the sgPFC (FDR<0.05) (**Fig. 1A**).

**Figure 1.**
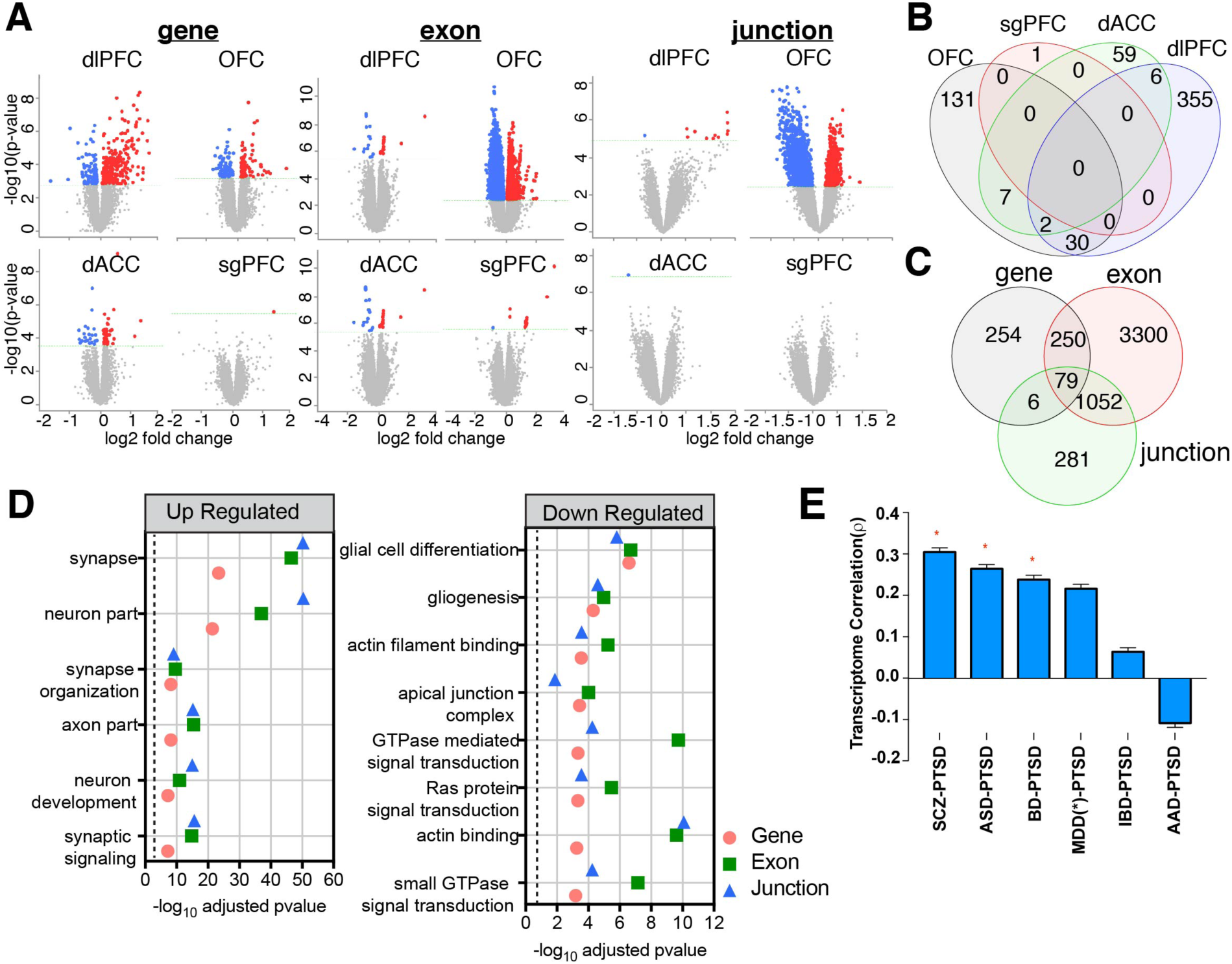
Global comparison of transcriptomic architectures across PFC subregions in PTSD and neurotypical controls. **a**, Volcano plots display differential regulation of three transcriptomic features: genes, exons, and exon-junctions for each cortical region. Blue dots indicate down regulation and red dots indicate up regulation compared to control. **b,** Venn diagram of overlap of PTSD DEGs in each cortical region. **c**, Venn diagram of overlap of differentially expressed gene features (genes, exons, and junctions). **d**, Top gene ontology (GO) enrichments for all differentially expressed features across the entire PFC including cell compartment, molecular function, and biological function ontologies. **e,** Rank ordered transcriptome similarity for all disease pairs with PTSD as measured with Spearman’s correlation of differential expression log_2_FC values.

We observed a small degree of overlap in the DEGs of each region, with the highest level in the dlPFC and OFC (30 DEGs) (**Fig. 1B**), lower levels between any 2 other regions, and only two genes (*ELK1* and *ADAMTS2*) that overlap across 3 regions (dlPFC, OFC, and dACC). No DEGs overlapped between all four regions. We observed significant overlap between gene features, especially between differentially expressed genes and exons and differentially expressed exons and junctions (**Fig. 1C**). Gene ontology (GO) analysis of all DEGs in the combined PFC using gProfiler (https://biit.cs.ut.ee/gprofiler/gost) revealed multiple GO terms, with the top terms plotted for all gene features (gene, exon, and junction) (**Fig. 1D; Table S2**); the top up-regulated gene ontologies included synapse, neuron, and axon terms, while the top down-regulated DEGs were associated with different aspects of glia formation, actin binding, and small GTPase signaling.

A recent large GWAS of PTSD subjects by the Psychiatric Genomics Consortium identified similarities in the genes with high polygenic risk scores (PRS) for PTSD with those with schizophrenia (SCZ) and bipolar disorder (BPD)^37^. This finding was also observed in the most recent PTSD GWAS published using the Million Veteran Program data set ^18^. Here we compared the PTSD transcriptomic signatures to a recent transcriptomic meta-analysis of other major psychiatric disorders, focusing on dlPFC ^38^. Our analysis of regionally specific transcriptomic correlations revealed similarly higher significant correlations of PTSD dlPFC gene expression with SCZ and BPD, as well as autism spectrum disorder (ASD), with a lower, nonsignificant correlation with MDD (**Fig. 1E; Fig S1**).

### PTSD transcriptomic organization differences revealed by network co-expression analysis

To characterize the transcriptomic changes in PTSD, we performed robust weighted gene co-expression network analysis (rWGCNA) ^39^ and identified 66 modules (arbitrary color designation) and plotted the modules with significant numbers of DEGs in **Fig. 2A** (see **Table S3** for complete list and membership). We employed ARACNE (Algorithm for the Reconstruction of Accurate Cellular Networks) to resolve the two-dimensional structure of the most significant individual modules and identify significant, highly interconnected key drivers that are differentially expressed and are predicted to drive observed transcriptional changes within modules ^40^. The module with the highest significance for association with PTSD, as well as number of DEGs and key drivers (coral2) (**Fig. 2B)** is composed of 36 genes, 7 of which are key drivers; coral2 is comprised primarily of downregulated DEGs in the dlPFC (FDR=5.9×10^-12^) and OFC (FDR=0.022). Notably, coral2 contains a number of down-regulated GABA-related DEGs (noted by green connections) in dlPFC that are also key drivers, including *GAD2* (glutamate decarboxylase 2), *SST* (somatostatin), *PNOC* (prepronociceptin), and *SLC32A* (solute carrier family 32, member 1). In addition, *ELFN1* (extracellular leucine rich repeat and fibronectin type III domain containing 10) and *LHX6* (LIM homeobox 6) are down-regulated key drivers that are linked with regulation of GABA synapse formation and GABA neuronal differentiation, respectively ^41–43^. We explored the possibility of transcriptome-wide changes to GABA related transcripts in PTSD (**Fig. 2C**) and observed significant decreases in transcripts encoding GABA transporters (SLC32A1) and GABA-related neuropeptides (*SST, PVALB, VIP, PNOC*) with concurrent increases in GABA receptors and anchoring proteins (*GABRA1, GABRA2, GPHN*) indicating a possible compensatory response to loss of GABA function (**Fig. 2C**). Network comparisons can provide identification of cell-type marker genes within modules and allow for identification of PTSD-associated networks within specific cell types. Coral2, as well as 5 other modules were significantly enriched for markers of neuronal cell type. Other top identified modules were significantly enriched for markers of endothelial cells (4 modules), oligodendrocytes (3), microglia (6), and astrocytes (4) (**Fig. S2**).

**Figure 2.**
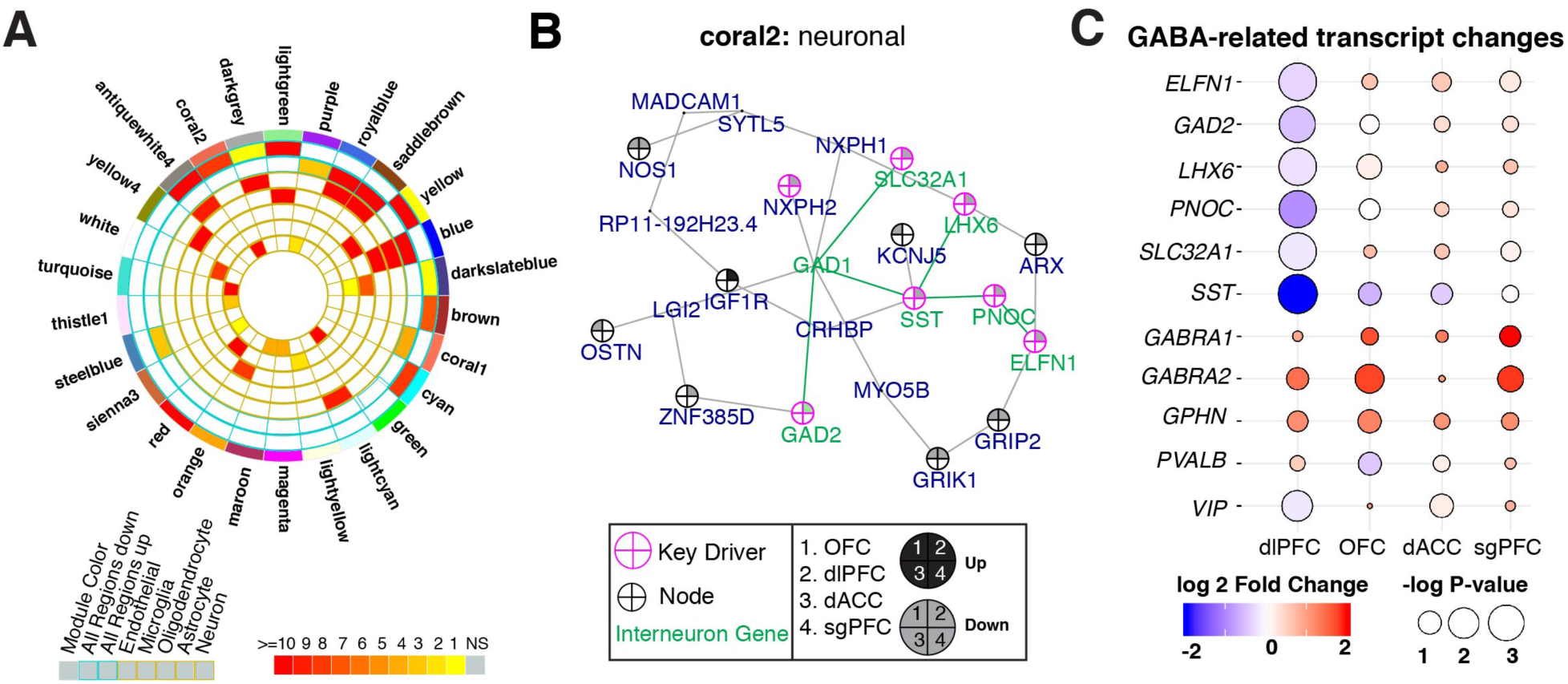
Regional gene co-expression analysis of PTSD PFC and Transcriptome Wide-Association Analysis. **a**, Degree of enrichment for DEGs (P<0.05) in modules for PTSD. Each slice of the chart represents a coexpression module assigned a random color name. The outer most concentric circles represent total DEGs across cortical region. The inner circles indicate cell-type specific markers for each module. Colors represent corrected FET P-values and a key is provided. **b**, Key driver and node representation of the most significant PTSD associated module, coral2. There is enrichment of neuronal markers within coral2. Key drivers are outlined in pink and nodes are outlined in black with inner colors representing regulation of gene expression (up regulation black, down regulation grey) across brain regions in PTSD as indicated. Interneuron related transcript connections highlighted in green. **c**, GABA-related key drivers and transcripts exhibiting global and regional up-regulation (red circles with black borders) or down-regulation (blue circles with black borders).

### Transcriptome-wide association studies identify gene *cis*-regulated expression in PTSD

Transcriptome-wide association studies (TWAS) were conducted to identify genes with expression that is significantly associated with PTSD using UTMOST (Unified Test for MOlecular Signatures) ^44^. TWAS has several advantages, including grouping of gene variants, reducing multiple comparisons, increasing power of association testing ^45, 46^, and enabling identification of potential transcriptomic effects caused by disease associated genetic variation. Convergence of gene expression changes in a PTSD dysregulated brain region with both illness state and genetic risk for illness would yield the most significant pathogenic markers of PTSD. We used the GTEx database ^47^ of tissue specific eQTLs (expression quantitative trait loci) to perform UTMOST-TWAS on the summary statistics of the largest PTSD GWAS recently published by the Million Veteran Program, PTSD Check List (PCL-17; n=186,689 individuals) ^20^, which identified 77 high-confidence risk genes for PTSD. Because there is a combination of races in our PTSD transcriptomic analysis, we used the PCL-17 trans-ancestral (combining European- and African-American) summary statistics for UTMOST-TWAS. Because there were significantly fewer females compared to males in the MVP cohort we used the combined-sex genotype data. We ran UTMOST combining PCL-17 and GTEx eQTLs from three different groups: PFC regions (includes dlPFC, dACC, and combined cortical tissue), CNS tissue (all brain tissue combined), and all non-CNS tissue. UTMOST identified 17 genes using the cortical tissue eQTLs (**Fig. 3A**) and 27 genes from the CNS tissue eQTLs (**Table S4**). Seven cortical TWAS hits were also DEGs in the current transcriptomic data set (*ELFN1*, *GJC1*, *LRRC37A*, *MAPT*, *MST1R*, *RBM6*, and *ZHX3*). We identified 46 unique genes across all three tissues by TWAS (**Fig. 3B**), with cortical tissue accounting for ∼30% of the TWAS signal (**Fig. 3C**). Moreover, we identified 17 DEGs that overlapped with the list of 77 high-confidence PTSD risk genes in the MVP study. Notably, we identified the highly connected GABA-related gene *ELFN1.* Further, our CNS TWAS identified the key driver *UBA7* (Ubiquitin like modifier activating enzyme 7) (**Fig. 3D**) as meeting genome-wide significance (**Table S4**).

**Figure 3.**
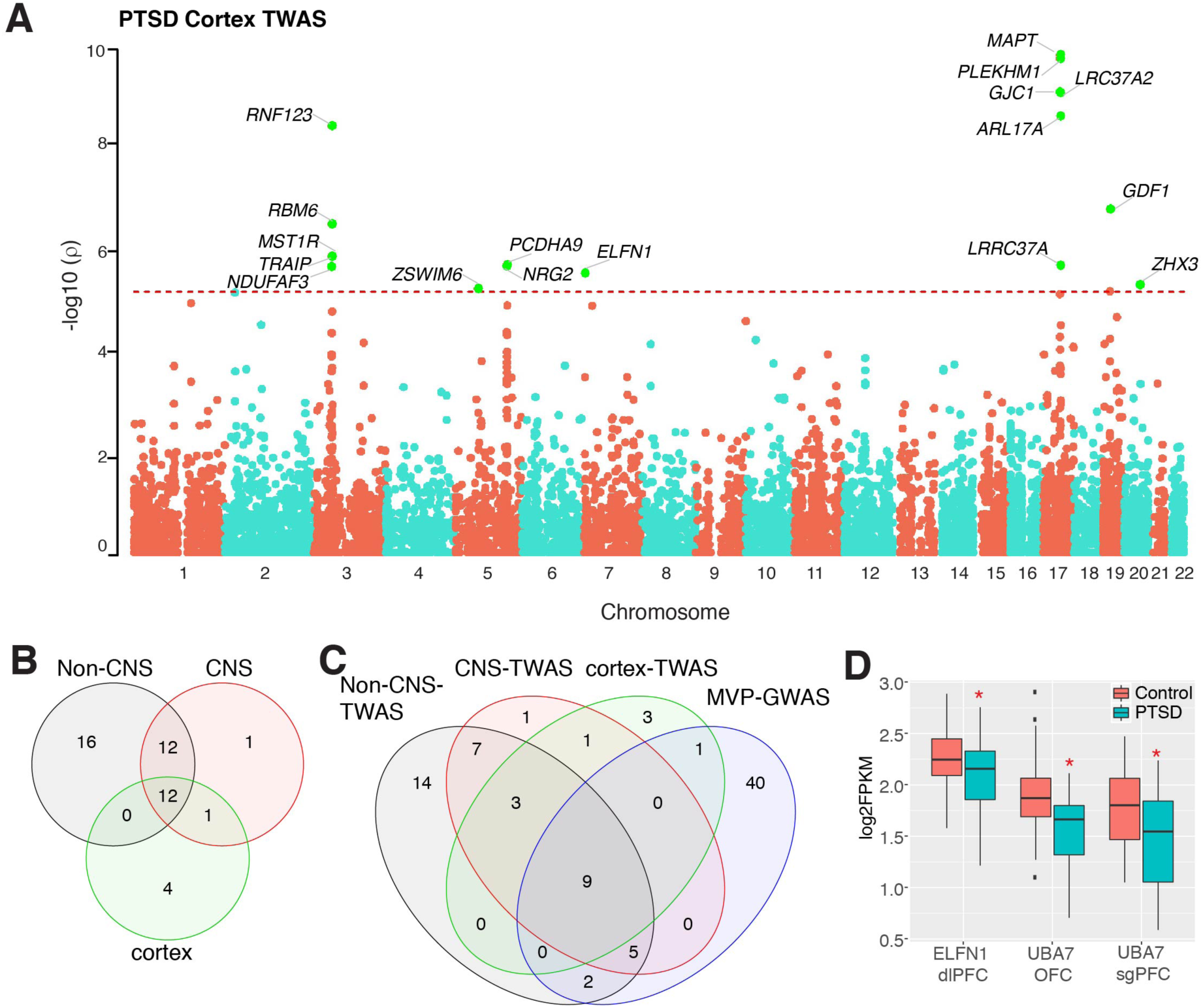
PTSD TWAS a, Manhattan plot of transcriptome-wide association by UTMOST using cortical tissue expression. Points above the red line indicate genes significant for PTSD. **b**, Venn diagram of gene overlap between TWAS tissue comparisons. **c**, Venn diagram of gene overlap between TWAS comparisons and MVP-GWAS. **d**, RNA-seq fold changes for ELFN1 in dlPFC and UBA7 in OFC and sgPFC. (*p<0.01, FDR<0.05, bars represent S.E.M. of FPKM).

### Sex-specific transcriptomic changes in PTSD

We observed significant principal component (PC) effects for primary diagnosis (Dx; PC 2, P=0.0003), sex (PC 1, P=2.2 ×10^-29^; PC 2, P= 2.3×10^-72^), age (PC 1, P=0.02), PMI (PC 1, P=0.0007), RIN (PC 1, P=1.5×10^-8^), and race (PC 1, P=0.01) on total variance across all brain regions (**Table S5**) and all of these factors were included in our model as covariates. Notably, sex had the most robust effect on variance with PTSD and there was a significant interaction for sex in our statistical model, indicating that there are highly significant sex-specific differences in the PTSD transcriptome.

Significant transcriptomic differences between males and females were observed in all PFC subregions (**Fig. 4A,B; Table S6**). Women exhibited more genes that were differentially expressed between PTSD and healthy controls: 2045 DEGs in the OFC and 1283 in the sgPFC, 38 DEGs in the dlPFC, and 0 in the dACC (**Fig. 4A; Table S6**). The relatively large number of DEGs in female sgPFC and OFC compared to the combined-sex data set (**Fig. 1A**) or male alone analysis (**Fig. 4B**) suggests that these regions are of particular importance to the pathophysiology of PTSD in women. Men exhibited many fewer DEGs comparing PTSD and controls, with the majority occurring in the dlPFC (84) and fewer in the OFC (2), sgPFC (2), and the dACC (1) (**Fig. 4B; Table S6**). There were very few regional DEGs overlapping between males and females (**Fig. 4C**).

**Figure 4.**
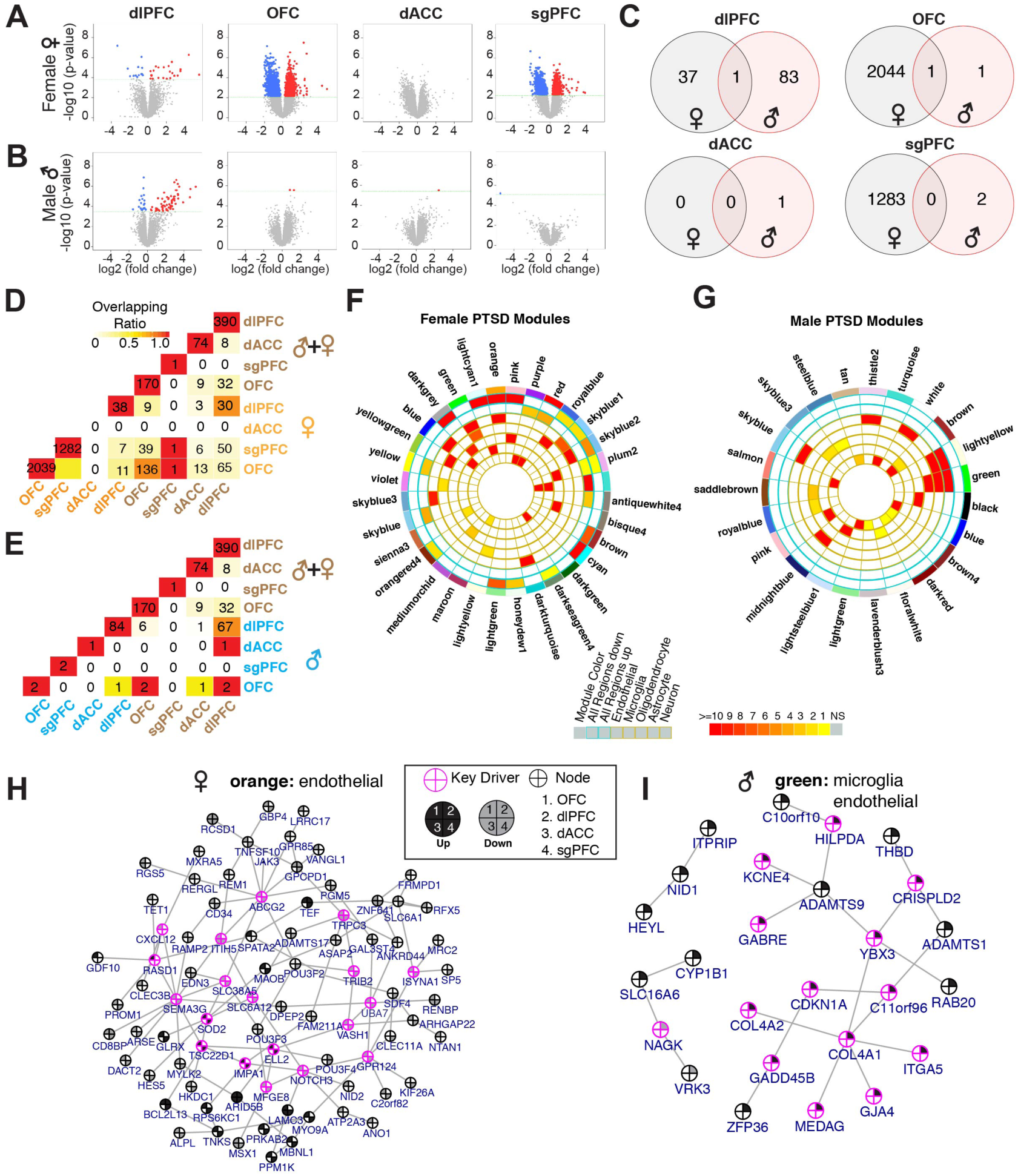
Differential expression profiles reveal distinct sex-specific PTSD transcriptomic signatures across PFC regions. Genes differentially regulated in female (**a**) and male (**b**) PTSD PFC regions. **c**, Regional overlap in the number of DEGs between females (black) and males (red). **d**, Regional overlap in number of DEGs compares combined sex (brown) and female only DEGs (orange). **e,** Compares combined sex (brown) and male only DEGs (blue). Lighter color indicates lower number of overlapping genes, with progressively darker red indicating increasing numbers of overlapping genes. Circos plots displaying the degree of enrichment for DEGs (P<0.05) in female (**f**) and male (**g**) modules for PTSD. Each slice of the chart represents a coexpression module assigned a random color name. The key for the concentric circles is shown in the middle. The outer most concentric circles represent total DEGs across cortical region. The inner circles indicate cell-type specific markers for each module. Colors represent corrected FET P-values and a key is provided. Key driver and node representation of the most significant sex-specific PTSD associated modules, orange in females (**h**), and green in males (**I**) enriched for DEGs for each comparison. There is enrichment of endothelial markers within orange, and endothelial and microglial markers within green. Key drivers are outlined in pink and nodes are outlined in black with inner colors representing regulation of gene expression (up regulation black, down regulation grey) across brain regions in PTSD.

Cross-regional comparisons revealed strong overlap between the female OFC and sgPFC (622 genes) (**Fig. 4D**), but little cross-regional overlap in males (**Fig. 4E**). Further analysis indicated that the small overlap observed between DEGs in males and females with PTSD is not due to baseline sex differences in gene expression (**Table S7**). These findings indicate that males and females with PTSD have unique, regional transcriptomic profiles with very little correlation in degree of transcript change or total number of DEGs.

### Sex-specific PTSD transcriptomic organization and key driver differences revealed by gene co-expression analysis

Male and female co-expression networks that combined all four PFC regions identified numerous transcript co-expression modules (**Fig. 4F and G**). In the female WGCNA, we identified 69 modules, 68 of which contained female-specific DEGs (**Fig. 5A** and **Table S3**) and in the male there were 59 modules, 18 of which contained male-specific DEGs (**Fig. 5B** and **Table S3**). We identified modules sharing high levels of homology at the gene membership level (**Fig. S3**; P<0.05; fold enrichment >20), notably, 43 of 69 female PTSD modules were conserved in male PTSD.

**Figure 5.**
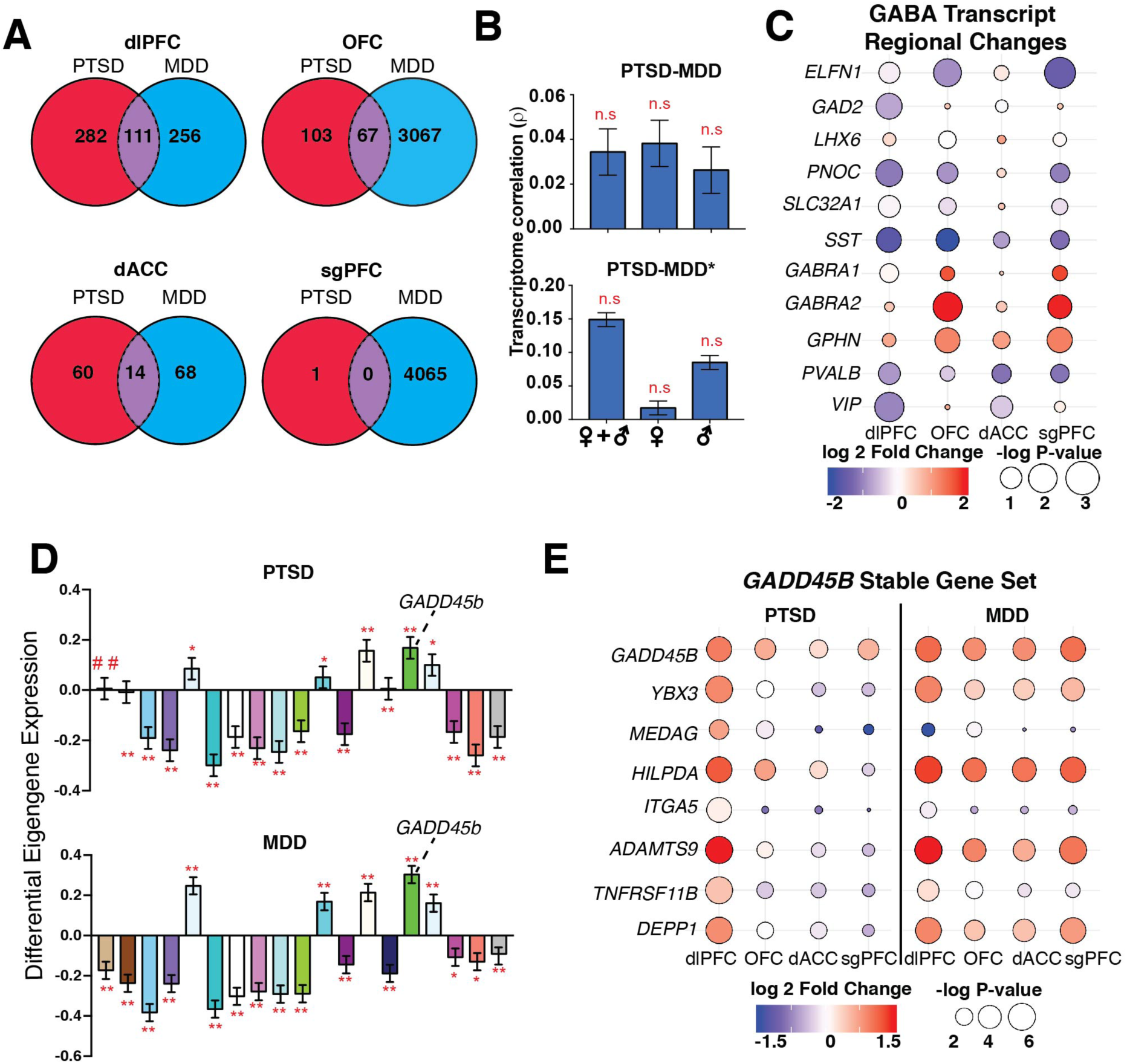
Gene and co-expression networks capture divergent and shared disease-specific features between PTSD and MDD. **a**, Venn diagrams displaying a low level of overlap in DEGs between PTSD (red) and MDD (blue) across PFC regions. **b,** Comparison of PTSD and MDD (top, current study) or meta-analyzed MDD profile (MDD*, from^38^), as measured with Spearman’s correlation of differential expression log_2_FC values showing no significant (n.s.) effects. **c**, Regional expression of GABA transcripts in MDD with up-regulation indicated by red and down-regulation indicated by blue circles. **d,** Module level differential expression across PTSD (top) and MDD (bottom). Plots show β values of module eigengene association with disease (FDR-corrected #P<0.1, *P<0.05, **P<0.01, error bars indicate SEM). **e,** A stable gene set that includes *GADD45b* is indicated on the module memberships in both disease conditions. This gene set is significantly stable (DEGs in both PTSD and MDD across all sex-specific comparisons) and indicates a likely point of molecular convergence between the two diseases, particularly in the dlPFC.

Examination of cell-type marker genes within modules revealed several sex specific PTSD modules that were enriched for cell-type marker genes; there were 7 modules enriched for endothelial cell markers in the female comparisons and 6 neuron-enriched modules in the male comparison, indicating a sex-specific divergence in cell-type associated molecular changes within the PFC (**Figure S2**).

The male and female PTSD co-expression modules containing DEGs were further examined for high network connection key drivers. The most significantly PTSD-associated female module (orange) is composed of 106 genes, 19 of which are key drivers, and is enriched with genes predominantly involved in immune function as well as endothelial cell markers (**Fig. 4H**); this module is composed of mostly downregulated genes across all PFC regions: sgPFC (FDR=2.29×10^-17^), dACC (FDR=3.47×10^-10^), OFC (FDR=7.27×10^-25^), and dlPFC (FDR=0.007; **Figure 4H**). In contrast, the most significantly associated male module (green), which contains 668 genes of which 14 are key drivers, is enriched for markers of both microglia and endothelial cell-types (**Fig. 4H**); DEGs in this module are predominantly upregulated in the dlPFC (FDR=3.58×10^-14^; **Fig. 4H**).

We validated the RNA-seq findings on a subset of genes identified as key drivers in our network co-expression analysis (**Table S3**). We performed quantitative real-time PCR on 39 individual key drivers identified in the top modules in figures 2 and 3. In the female comparison 90% of the key drivers (28 of 31) were significantly replicated in the correct, up or down direction; in males 100% (12 of 12) and in the combined-sex comparison 90% (9 of 10) of key driver DEGS replicated in the correct direction PCR (**Table S8**).

### Divergent and convergent genomic mechanisms between PTSD and MDD

The MDD psychiatric control group was assembled to control for quality of the tissue, manner of death (number of suicide cases-a known confounder), antemortem treatment, and substance abuse/dependence, as well as the high rate of comorbidity with PTSD (approximately 50% of newly diagnosed PTSD patients) ^48^. We performed RNA-seq and bioinformatics analyses on the MDD cohort in the same manner as the PTSD cohort (summarized in **Figure S4 and Tables S9** and **S10)**. The greatest numbers of MDD DEGs were observed in the sgPFC (4065) and OFC (3067), regions with relatively few DEGs in the PTSD cohort (Figure 5a). Further analysis revealed moderate to low levels of overlap between MDD and PTSD DEGs; 111 in dlPFC, 67 in OFC, 14 in dACC, and no overlap in sgPFC (**Fig. 5A**). To further explore the transcriptomic relationships between PTSD and MDD, we compared the differentially expressed log2 fold changes of the combined cortical regions and found very low (ρ<0.05), nonsignificant correlations between the sex-combined, female, or male analyses (PTSD vs. MDD, **Fig. 5B**). To extend and validate this finding the same comparison was performed using a recently published, aggregated, meta-analyzed MDD* profile (PTSD-MDD*)^38^, which shows higher (ρ<0.15), but still nonsignificant correlations. These findings are consistent with the disease correlation analysis in **Figure 1E**, showing that the PTSD transcriptome significantly correlates with SCZ, BPD, and ASD, but not MDD*. Similar to previous reports ^30, 49^, we also observed sex-specific transcriptomic signatures in MDD (**Fig. S4B & C**, **Table S10**). Since GABA related deficits have been reported in MDD ^50, 51^ and to compare with the current findings of GABA alterations in PTSD (**Fig. 2B, C**), we examined GABA signaling transcripts in our MDD data set. We found significant GABA-related DEGs in MDD, some of which overlapped with PTSD, particularly in the dlPFC (e.g., *GAD2, SST, PNOC, VIP*) (**Fig. 5C**); there were also dlPFC differences, with *PVALB* levels decreased in MDD and a small non-significant increases in PTSD. There were also regional differences between the OFC and sgPFC where we observed decreases in *ELFN1, PNOC*, and *SST* in MDD but no effects in PTSD (**Fig. 5C**)

The disparity in molecular signatures between PTSD and MDD was also tested at the level of network analysis. We calculated the correlation between MDD and PTSD for each module and the quantile of the correlation among 2000 permutations. Of the 47 combined sex MDD modules, 16 were convergent and 12 were divergent compared to PTSD (there was no effect on the remaining 19 modules); of the 59 female modules 9 are convergent and 19 are divergent; and of the 43 male modules 15 are convergent and 5 are divergent (**Fig. S5**). These findings indicate that, even for modules conserved on the gene membership level, relatively small subsets (∼10% for female modules, ∼35% for combined sex and male modules) display significant convergence between MDD and PTSD; notably, there were similar levels of divergent modules (∼12-25% for male and combined sex, 33% for female modules).

Due to the high comorbidity between PTSD and MDD and interest in identifying converging molecular pathways in neuropsychiatric disorders, we further explored the modules with high convergence between MDD and PTSD to identify co-expression gene sets common to both disorders across all PFC subregions. We calculated module-level differential expression across both disease states (**Fig. 5D**) and identified 19 modules with high correlation for PTSD and MDD in the combined-sex comparison, 18 in the female, and 6 in the male comparisons (**Fig S6A**) Of particular interest was the appearance of a stable gene set with high pair-wise connectivity present across all three comparisons, made up of 8 genes, including the DNA methyltransfease *GADD45B* (growth arrest and DNA damage induced 45b; **Fig. 5E**). We also observed cell-type specific marker enrichment in many of the PTSD-MDD converging modules, in astrocytes, microglia and oligodendrocytes (**Fig. S6B-D**).

## DISCUSSION

Here we present the first complete transcriptomic evaluation of human PFC in PTSD, made possible by the establishment of the National PTSD Brain Bank by the U.S. Department of Veterans Affairs ^52^. This is the largest transcriptomic study of PTSD brain tissue to date and concurrently examines the transcriptomic state for a psychiatric control, MDD, providing both a comparison and control for these comorbid illnesses. This is also one of the largest MDD transcriptomic data sets and includes regions (OFC) not typically examined ^53^. Further, we take advantage of data from the largest PTSD GWAS from the Million Veteran Program ^20^ to link risk loci and transcriptomic changes in PTSD.

Integration of transcriptomic data with genetic variant information enabled us to identify genes associated with PTSD risk and illness state. TWAS-UTMOST of CNS and cortical tissue identified 45 genes achieving genome-wide significance using the MVP data, two of which, *ELFN1* and *UBA7*, are also DEGs and key drivers with multiple network connections. *ELFN1* is downregulated in the dlPFC and is a key driver in the most significant combined-sex network (**Fig. 2B & C**); it is also decreased in female dlPFC and dACC and is a key driver in a female-specific network (**Table S3**). *ELFN1* encodes a synaptic adhesion protein that acts as an allosteric modulator of group III metabotropic glutamate receptors ^41^; it is also required for synapse formation on SST GABA interneurons by recruiting presynaptic mGluR7 (metabotropic glutamate receptor 7) and GluK2 (glutamate receptor kainate 2) ^42^. Notably, there are multiple GABA related transcripts that are significantly down-regulated key-drivers in the same network as *ELFN1*, including *SST*, *PNOC* (Prepronociceptin, a GABA interneuron maker), *GAD2* (glutamic acid decarboxylase-2, GABA synthetic enzyme), and *SLC32A1* (solute carrier family 32A1, GABA/glycine vesicular transporter), indicating a decrease in GABA interneuron transmission in PTSD (**Fig. 2C**). These effects were specific to the dlPFC where there was some overlap with MDD, as previously reported ^50, 51^. However, there were also significant differences in dlPFC (i.e., PVALB), as well as in OFC and sgPFC, with decreased GABA-related transcripts in MDD, but small increases or no effects in PTSD. The region-specific overlap in GABA transcripts in the dlPFC could account in part for PTSD and MDD comorbidity, while the differences in dlPFC, OFC and sgPFC could represent illness specific changes.

SST and other GABA interneuron deficits, identified in postmortem studies of MDD ^50, 51^ and SCZ ^54^, interfere with PFC gamma oscillations and thereby alter top-down inhibition of projection target regions ^55^. The GABA deficit overlap suggests that similar effects could occur in PTSD. This is supported by preclinical ^56–60^ and human brain imaging ^61, 62^ studies reporting that PTSD pathophysiology is also linked to PFC dysfunction, which could impair inhibitory control of the amygdala, as well as other target regions that regulate fear and extinction behaviors. Identification of significant links between *ELFN1* risk loci and decreased *ELFN1* expression, and its role as a key driver in the top PTSD network that includes GABA-related key drivers (*SST, PNOC, GAD2*) that are also decreased, provides strong support for a connection with PFC dysfunction in PTSD. However, differences between MDD and PTSD in PFC subregions demonstrate illness specific effects that could contribute to disruption of local PFC circuits as well as target regions. Based on these findings, future studies using high resolution brain imaging approaches could probe for these PTSD vs. MDD subregion specific effects.

*UBA7* is a key driver in a combined sex network (brown) (**Table S3**) and in the most significant female PTSD module (orange), which is also enriched for endothelial cell specific markers (**Fig. 4H**). *UBA7* is downregulated in the OFC in both comparisons and in female sgPFC. Transcriptomic studies of peripheral tissue implicate inflammatory and GC signaling in PTSD ^31^ and *UBA7* is a ubiquitin-like activating enzyme that is part of an inflammatory gene expression cascade ^63^. In addition, the glucocorticoid receptor chaperone *FKBP5* was identified as an upregulated key driver in the dlPFC of females with PTSD. Regulation of *FKBP5* has also been reported in small, preliminary postmortem PTSD studies in OFC ^64^ and the sgPFC ^65^ and *FKBP5* variants have been extensively implicated in risk for developing PTSD ^66, 67^. These results indicate that *FKBP5* and *UBA7* act as hub genes with functional importance in PTSD molecular pathology, consistent with previous work implicating dysregulation of inflammatory and GC signaling as hallmarks of PTSD pathophysiology. These targets regulate high confidence networks of genes, and in the case of *UBA7* are related to both risk loci and illness state and offer potential for therapeutic interventions.

Our results also demonstrate sex specific transcriptomic signatures across the PFC subregions, with relatively large numbers of genes regulated in most PFC regions of females, and fewer DEGs in males, except in dlPFC. Sex-specific, co-expression modules with only modest homology between females and male were also identified. These sex differences extend into the cell-type specificity of significant modules with female modules associated primarily with endothelial cells and male modules associated more with neurons. While the overall transcriptomic signature of male PTSD is markedly different from MDD (**Fig. 5 B**), this finding suggests male PTSD molecular pathology is more similar to depression than its female counterpart.

Recent evidence points to converging molecular mechanisms across neuropsychiatric disorders^38, 68, 69^, and because of the high rates of PTSD and MDD comorbidity (> 50%) we reasoned that MDD was an optimal psychiatric control to disentangle the differences and similarities in the two disorders. Consistent with previous reports ^30^, large numbers of DEGs were identified in all PFC subregions of both males and females with MDD, particularly sgPFC and OFC (**Table S10**) but there was little overlap with PTSD. We further tested the concordance of the complete PTSD and MDD transcriptomes identified in the current study, and with a previous MDD meta-analysis ^38^ but found no significant correlations (**Fig. 5B**). At the network level there were low levels of convergence, but there was a set of related genes that were consistently regulated across the combined-sex, female, and male PTSD and MDD modules. The overlap in this set of genes, which included *GADD45B*, was observed in the dlPFC, but there were also significant effects in the other subregions in MDD, but not PTSD. *GADD45B* is a stress and activity regulated DNA methyltransfease that has been implicated in long-term synaptic plasticity and cellular stress responses, and is a key modulator of epigenetic switching in neurological and psychiatric disorders ^70^. *GADD45B* functions in part via regulation of the promoters of *FGF1* (fibroblast growth factor 1) and *BDNF* (brain derived neurotrophic factor)^70^, and has been implicated in preclinical models of depression ^71^. There are currently no studies examining the role of *GADD45B* in preclinical models of PTSD, but the results indicate that this gene and its network partners represent a potential molecular intersection between PTSD and MDD pathology.

In conclusion, we present the first well-powered transcriptomic study of human postmortem brain, focused on PFC in PTSD and define molecular signatures that are common to both female and male PTSD subjects, as well as sex-specific transcriptional changes, including differentially expressed key drivers that are linked to PTSD genetic risk. The results provide strong evidence for the role of these and other key drivers in PTSD that will guide future functional studies in preclinical models. In addition, this extensive dataset represents an important resource and significant step forward in understanding the genomic architecture of the PTSD, as well as MDD brain by identifying the molecular pathways which are significantly involved in mediating the effects of trauma. Future investigations should include integrating brain transcriptomic profiles with those of peripheral tissue datasets to identify biomarkers of PTSD ^72^, and neuroimaging to characterize cortical differences between PTSD and MDD. The results of this study further elucidate the molecular mechanisms underlying PTSD and will lead to improved diagnostic criteria and therapeutic interventions.

## MATERIALS AND METHODS

### Data Reporting

No statistical methods were used to predetermine sample size.

### Human Subjects

Human autopsy brain tissue samples were obtained from the University of Pittsburg Medical Center and the National Center for PTSD Brain Bank (with consent of next of kin). Subjects were a mix of European-, Asian-, and African-American descent. Males and females were group-matched for age, pH, and postmortem interval (PMI).

Sociodemographic and clinical details are listed in **Table S11** and includes presence of comorbid disorders, tobacco use, cause and manner of death and presence of drug and/or alcohol abuse. Inclusion criteria for PTSD, MDD and control cases were as follows: PMI<30 hours, age range >18 to <70. One hundred and forty-three subjects (52 PTSD: 26 males, 26 females; 45 MDD: 27 males, 18 females; and 46 controls (CON): 26 males, 20 females) were used in this study. Fresh frozen samples (50mg) of prefrontal cortex from subgenual prefrontal cortex (Brodmann area 25), dorsal anterior cingulate cortex (Brodmann area 24), medial orbitofrontal cortex (Brodmann area 11), and dorsolateral prefrontal cortex (Brodmann area 9/46) were collected from each postmortem sample.

Psychiatric history and demographic information were obtained by psychological autopsies performed postmortem. Trained clinicians with informants best acquainted with the subject generated the necessary information for diagnosis. To avoid systematic biases PTSD, MDD and CON cases were characterized by the same psychological methods. Psychiatric diagnosis was determined by consensus diagnosis between two clinicians using DSM-IV^73^ criteria using SCID-1 interviews that are adapted for psychological autopsy^74^ and review of medical records.

### RNA-sequencing and Bioinformatics

RNA (20mg) from each PFC region was extracted using RNeasy Mini Kit with gDNA elimination, as described by the manufacturer (Qiagen). RNA integrity number (RIN) and concentration were assessed using a Bioanalyzer (Agilent). Libraries were constructed using the SMARTer® Stranded RNA-seq Kit (Takara Bio) preceeded by rRNA depletion using 1ug of RNA. Samples were bar coded for multiplexing and sequenced at 75 bp paired-end on an Illumina HiSeq4000. Samples were pooled eight per lane and sequenced at a depth of 50 million reads.

#### Data preprocessing and quality control

Sequences in FASTQ files were mapped to the human genome (Hg19) using *STAR* (version 2.5.3a) with reference genome and annotation GTF file downloaded from ENSEMBL (release 79, GRCh38), and then counted by *featureCounts* (version 1.5.3). ENSEMBL IDs were mapped with gene annotation using the *biomaRt* package in R. Samples with overexpressed mitochondria genes that account for more than half of total counts were filtered out.

#### Differential gene expression (DGE)

DGE was calculated using the *DESeq2* package in R for each sex and brain-region specific group^75^. The statistical framework enabled calculation of log2 fold-change values (log_2_FC) for each gene and disease from raw counts. This model takes into account covariates including age at time of death, race, PMI (post-mortem interval), and RIN (RNA integrity number). Genes with zero expression in all samples in a group are dropped. Differential expressed genes are defined by a cut-off adjusted P value(*P*<0.5) and an FDR < 0.05 (Benjamini-Hochberg). We employed an adapted protocol ^76^ on BAM files to extract transcript feature counts from each cortical subregion using regtools^77^ and bed_to_junctions from TopHat2^78^.

#### Gene co-expression network analysis

Network analysis was performed with the *WGCNA* package using signed networks, either in sex and brain-region specific groups or in sex and disease-specific groups^39, 79^. Data was quantile normalized for GC-content using R package *cqn* and corrected for batch effects using *ComBat* by package *sva*. Outliers were defined as samples with standardized sample network connectivity Z scores < −2 and were removed. A soft-threshold power of 6 was used for all studies to achieve approximate scale-free topology (R^2^>0.8). Networks were constructed using the *blockwiseModules* function. The network dendrogram was created using average linkage hierarchical clustering of the topological overlap dissimilarity matrix (1-TOM). Modules were defined as branches of the dendrogram using the hybrid dynamic tree-cutting method, with a minimum module size of 20, merge threshold of 0.1, and negative pamStage. Modules are labelled by a random color for illustration purposes. Genes that did not fall within a specific module are assigned the color grey. Module (eigengene)-disease associations were evaluated using a linear mixed effect model. We also used linear regression to test for association between module eigengenes and several covariates or confounders (age, race, PMI etc). Significance values were FDR-corrected to account for multiple comparisons. Genes within each module were prioritized based on their module membership (k_ME_), defined as correlation to the module eigengene.

#### Transcript Network Organization Analysis

After retrieving log2 fold changes of each transcript in different sexes by region combinations, we assigned a significance level to each gene in modules defined by WGCNA modular analysis pipeline. The WGCNA analysis generated 4 background networks with different total modules that included female control, female PTSD, male control and male PTSD. Each module was named with a color which would be used for visualization and identification. Background networks were created to integrate differentially expressed genes (DEGs) and modular changes from different regions. The number of identified DEGs was counted for each module and a *Fisher’*s exact test was utilized to test if DEGs are enriched in a specific module. The network structures of significant modules were visualized using *igraph* in R. Key drivers which are defined by connectivity with DEGs were labeled in network graphs. These analyses were done for modules in each background network.

To further identify key regulator (driver) genes of the modules identified by WGCNA, we applied the key driver analysis to the module-based weighted coexpression networks derived from WGCNA. WGCNA was used first to identify significant interactions between genes in each module based on their mutual information and then remove indirect interactions through data processing. An undirected Key Driver Analysis was performed using the R package *KDA*. *KDA* calculates connectivity to a specified depth between each signature gene and a hub signature gene with connection to a significant number of other signature transcripts defined as a key driver.

#### Correlation/Concordance with other Psychiatric disorders

Microarray and RNA-seq data from 700 postmortem cortical brain samples across 14 studies of multiple neuropsychiatric disorders (including MDD) were meta-analyzed in a previous study^38^. We used Spearman’s ρ to compare those DEG meta-analysis log_2_FC signatures to the PTSD and MDD DEG log_2_FCs in our cohorts and compared for all disease pairs.

#### UTMOST-TWAS

UTMOST (Unified Test for MOlecular SignaTures) is a principled method to perform cross-tissue expression imputation and gene-level association analysis and was used to identify PTSD-associated genes based on GWAS results from the PGC and MVP. It takes in SNP-level summary statistics and calculates the probability of each gene to be associated with PTSD in a specific tissue or jointly analyzed across multiple tissues based on tissue-specific eQTL information. The P-values are than adjusted by Bonferroni correction and genes with adjusted P values < 0.05 are reported as meeting genome-wide significance. PGC-PTSD sex-and-race-specific or sex-specific GWAS summary statistics are calculated by UTMOST using eQTLs from cortical tissues, CNS or non-CNS databases. The most current version of UTMOST uses GTEx Analysis release V6p and full list of 44 tissues can be found on GTEx website (Genotype-Tissue Expression, https://gtexportal.org/home/).

### Quantitative PCR Methods

qRT-PCR was performed using primers designed to detect the transcript of all 40 key drivers in the most significant co-expression modules for the combined sex, female, and male. mRNA was isolated from the using the RNEasy Plus Mini Kit (Qiagen, Venlo, Netherlands); 1 ug of mRNA was reverse-transcribed into cDNA using the iScript cDNA Synthesis kit (Bio-Rad, Hercules, CA). RNA was hydrolyzed and re-suspended in nuclease free water. Gene specific primers for were designed using Primer 3 v.0.4.0 freeware (http://bioinfo.ut.ee/primer3-0.4.0/) and tested for efficiency and specificity by serial dilution and melt curve analysis. Sybr Green mix (Bio-Rad, Hercules, CA) was used to amplify cDNA. Fold regulation was calculated by using the –delta delta Ct (2-DDCt) method. The 2-DDCt analysis calculates relative gene expression levels between two samples by using the threshold cycles calculated by increasing fluorescent signal of the amplicon during PCR.

### Code Availability

All code used in this study is freely available online and can be found at https://github.com/mjgirgenti/PTSDCorticalTranscriptomics

### Data Availabilit

RNA-seq summary statistics generated and/or analyzed during the current study will be made available on GEO. Additionally, the data that supports the findings of this study are available for from the corresponding authors upon reasonable request.

## CONSORTIA

Names of all consortium members are available from the authors upon request.

## ACKNOWLEDGEMENTS

We would like to thank Dr. Marina Picciotto and Dr. Christopher Pittenger for critically reading the manuscript. We would like to thank Rosemarie Terwilliger for technical assistance. This work was supported with resources and use of facilities at the VA Connecticut Health Care System, West Haven, CT, Central Texas Veterans Health Care System, Temple, TX, Durham VA Healthcare System, Durham NC, VA San Diego Healthcare System, La Jolla, CA,, VA Boston Healthcare System, Boston, MA, USA and the National Center for PTSD, U.S. Department of Veterans Affairs. The research reported here was supported by the Department of Veterans Affairs, Veteran Health Administration, VISN1 Career Development Award to M.J.G. and by NIMH grants MH093897 and MH105910 to R.S.D.

## AUTHOR CONTRIBUTION

M.J.G., M.J.F., J.H.K., and R.S.D conceived the project, designed the experiments and wrote the manuscript. M.J.G. also generated and analyzed all of the data. J.W., D.J., H.Z. oversaw all bioinformatics analyses for gene expression, network analysis, and TWAS. M.B.S. and J.G. contributed GWAS data. D.C., K.Y., B.R.H., D.E.W. contributed to study design. All authors contributed to the preparation of the manuscript.

## DISCLAIMER

*The views expressed here are those of the authors and do not necessarily reflect the position or policy of the Department of Veterans Affairs (VA) or the United States government*.

---

**Supplementary Figure 1.**
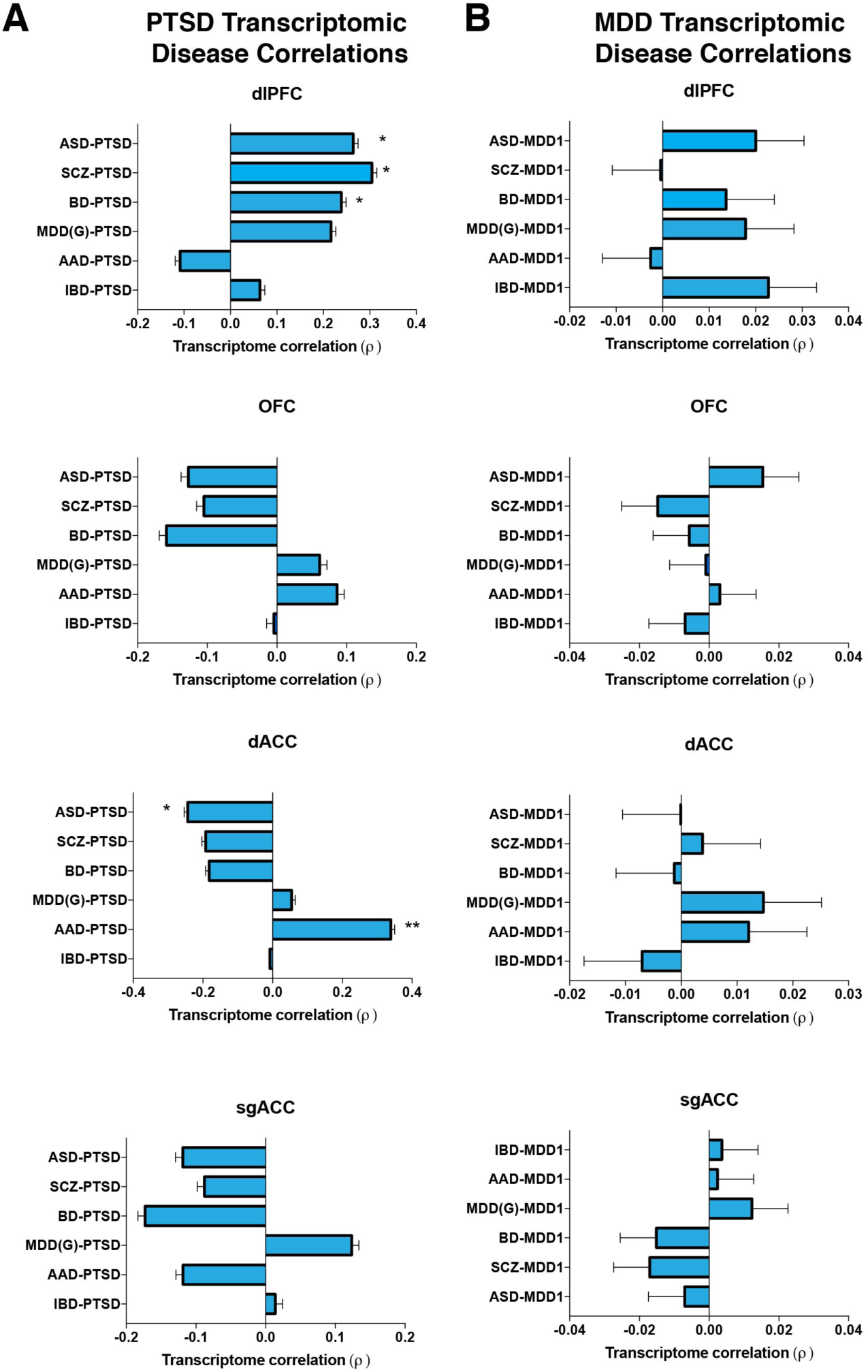
Transcriptomic concordance comparisons between PTSD and other neuropsychiatric disorders. **a**, PTSD transcriptomic profiles were compared with 5 other neuropsychiatric disorders. Individual cortical regions were compared to meta-analyzed transcriptomic signatures for autism spectrum disorder (ASD), schizophrenia (SCZ), bipolar disorder (BP), major depression disorder (MDD*), alcohol abuse disorder (AAD), and control disease irritable bowel syndrome (IBS). There was a significant correlation between PTSD and ASD, SCZ, and BP in the dlPFC and significant correlation between PTSD and AAD in the dACC. **b**, Comparison of psychiatric control cohort MDD transcriptomic signature to the 5 psychiatric disorders.

**Supplementary Figure 2.**
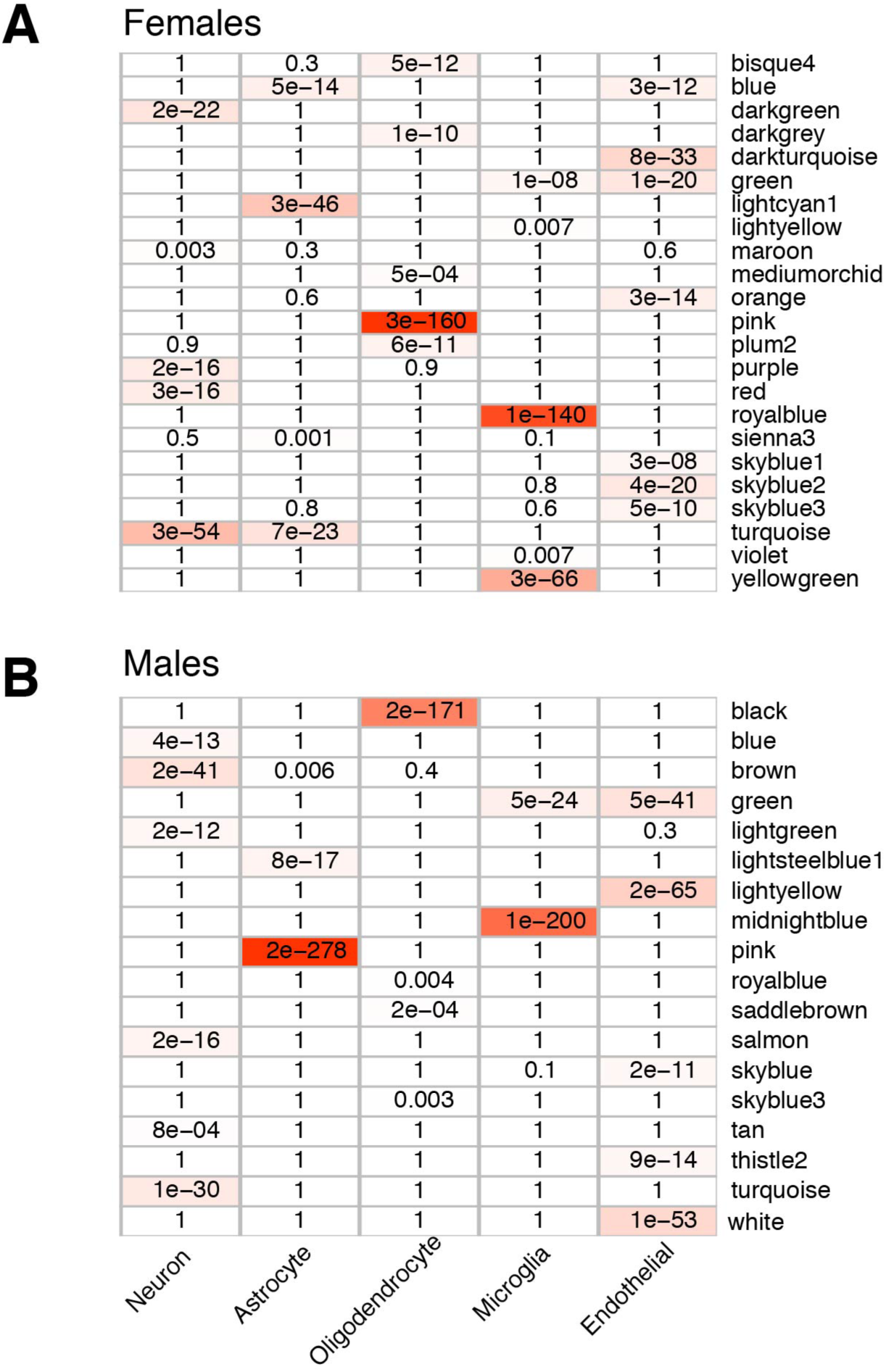
Sex-specific PTSD associated modules with enrichment of cell-type markers. Cell type enrichment was found in 23 female specific modules (**a**), and 18 male specific modules (**b**).

**Supplementary Figure 3.**
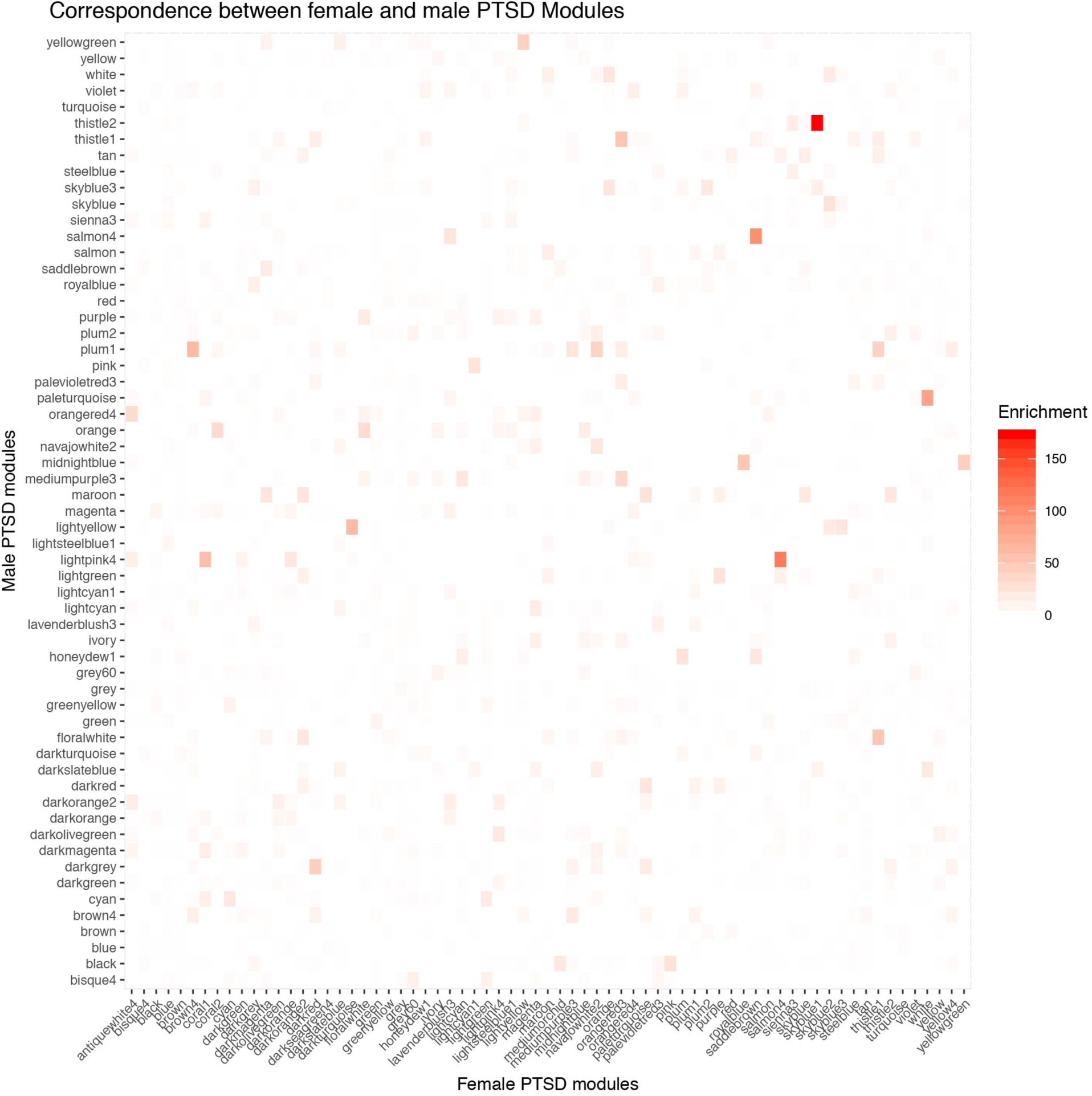
Correspondence between female and male PTSD coe-expression modules. Homology was measured between all male and female modules with significant association with PTSD. Significant homology was observed in 42 modules with greater than 20% enrichment in module gene membership between females and males.

**Supplementary Figure 4.**
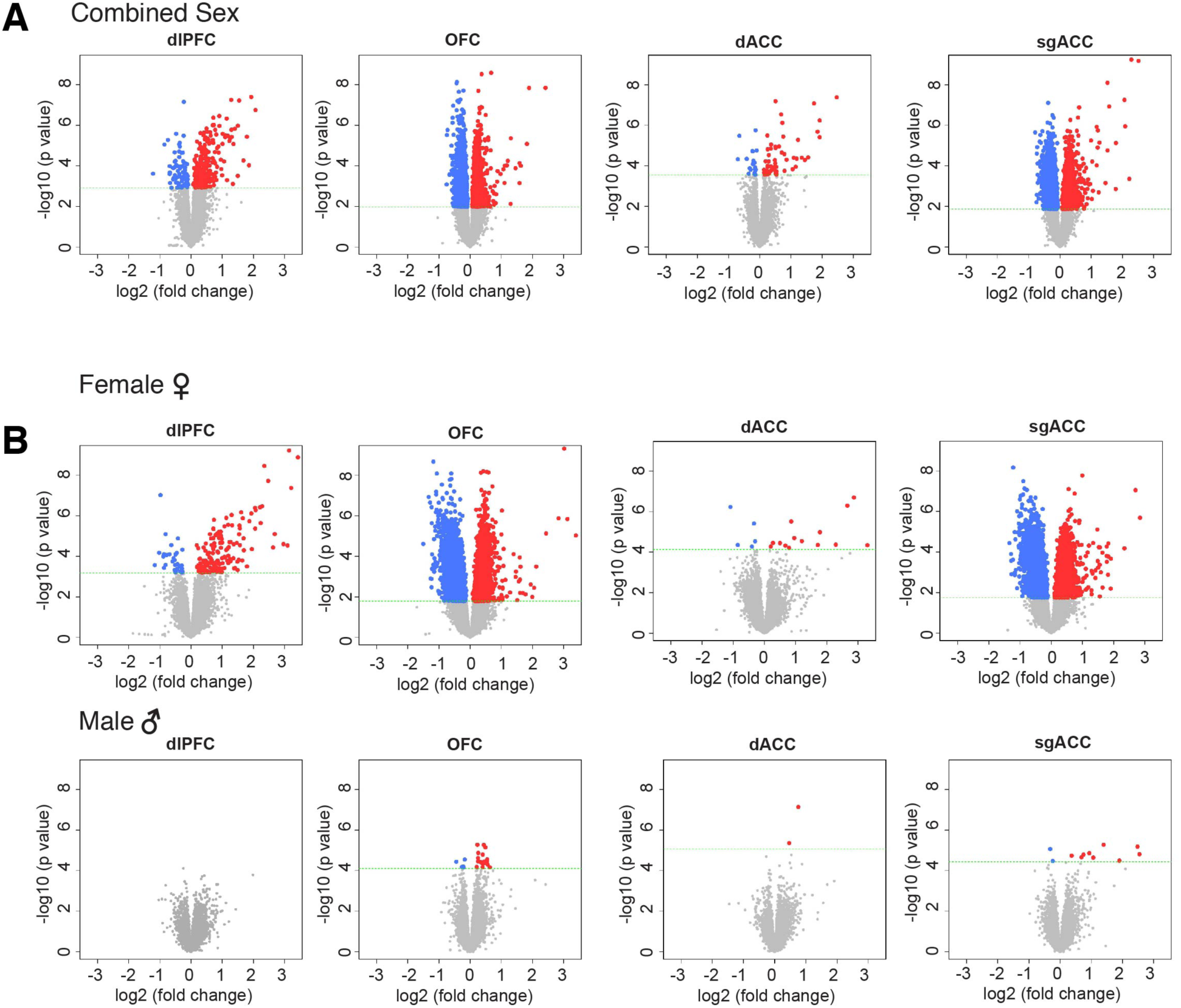
Differential expression profiles in humans with MDD reveal sex-specific transcriptomic profiles across PFC subregions. **a**, Volcano plots display differential regulation of genes in PFC of all MDD cases. **b**, Volcano plots display sex-specific DEGs in females (top) and males (bottom) across PFC subregions. Blue dots indicate down regulation and red dots indicate up regulation compared to control.

**Supplementary Figure 5.**
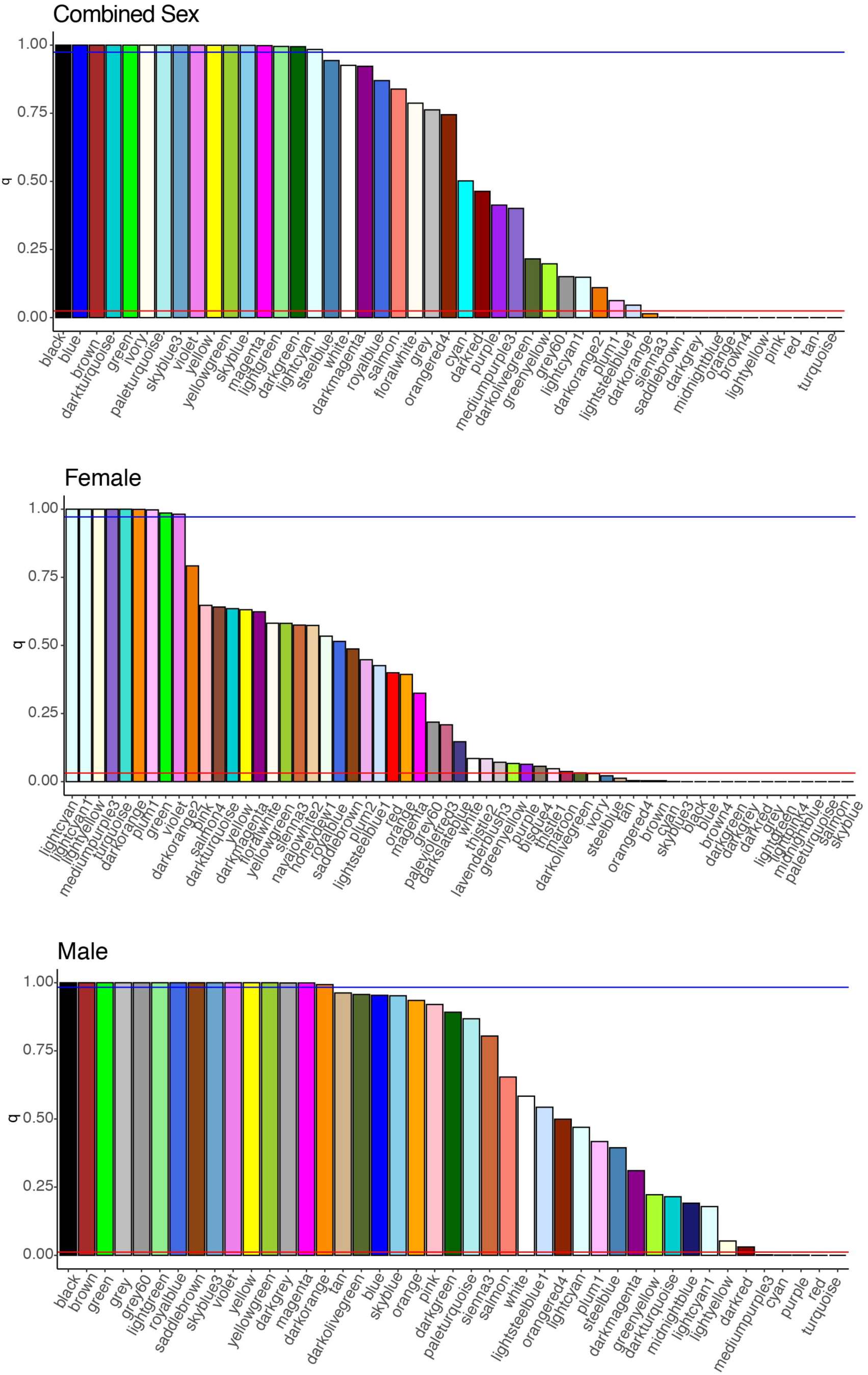
Module correlation between PTSD and MDD gene co-expression modules. The quantile of correlation among 2000 permutations was calculated for each PTSD and MDD module. Blue line indicates modules above the 97.5% quantile that are significantly convergent and the red line indicates the bottom 2.5% quantile that are significantly divergent. In the combined sex comparison(**a**) there are 16 convergent modules and 12 divergent modules, in females there are 9 convergent and 19 divergent modules(**b**), and in males there are 15 convergent and 5 divergent modules(**c**).

**Supplementary Figure 6.**
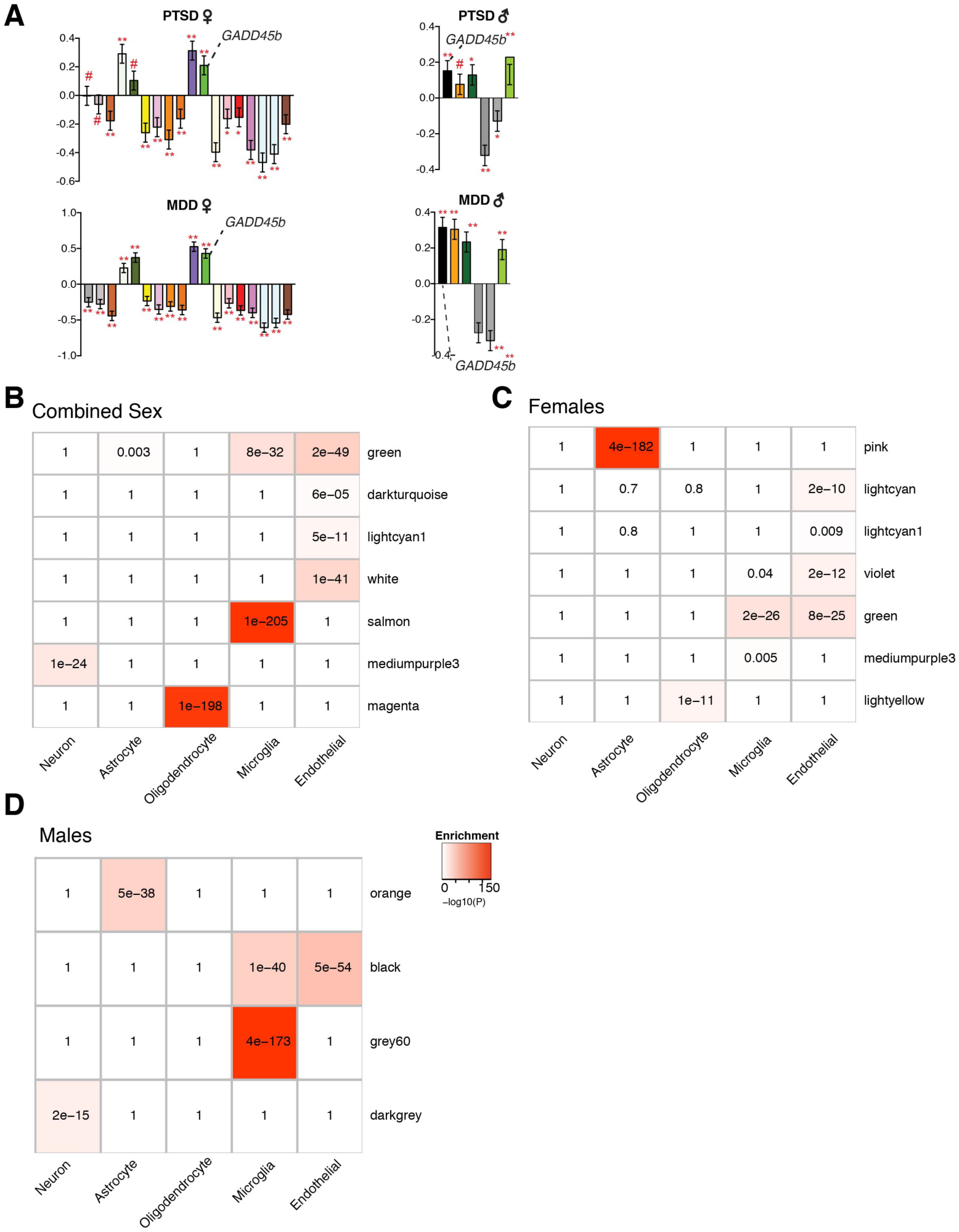
Sex- and cell-type specific network convergence of PTSD and MDD. **a,** Female and male module level differential expression across PTSD (top) and MDD (bottom). Plots show β values of module eigengene association with disease. FDR-corrected #P<0.1, *P<0.05, **P<0.01, error bars indicate SEM). Cell type marker enrichment of combined sex (**b**), female (**c**) and male (**d**) modules in combined disease network analysis.

**Supplementary Figure 7.**
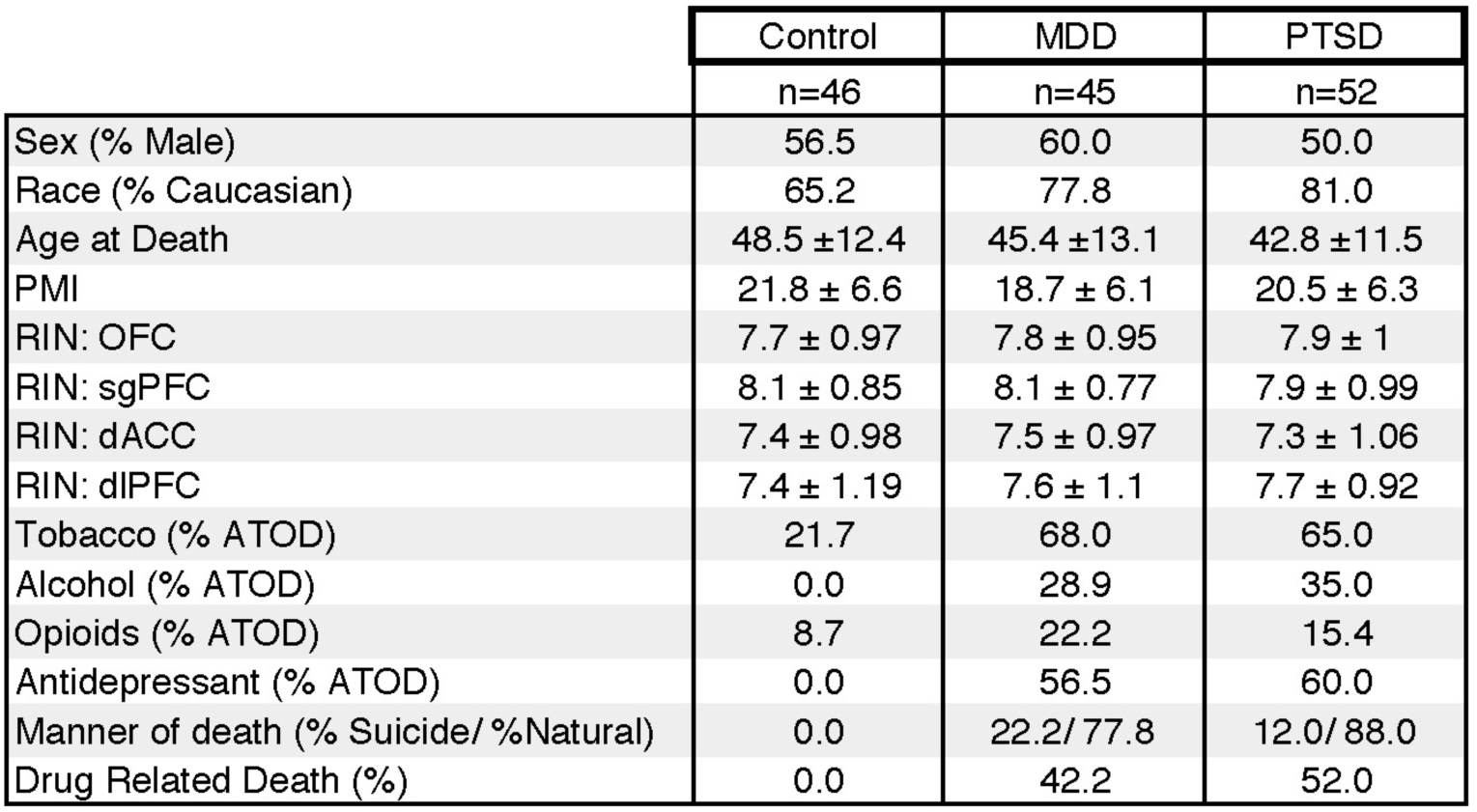
Sociodemographic and clinical summary of Control, MDD, and PTSD cohorts.

**Supplementary Table 1.** List of genes differentially expressed across brain regions in PTSD vs. control.

**Supplementary Table 2.** Summary of gene ontology analyses performed on PTSD DEGs.

**Supplementary Table 3.** Summary Table of gene network analysis results in combined sex, female and male PTSD.

**Supplementary Table 4.** Summary of significant TWAS hits for cortical, CNS, and non-CNS analyses using MVP GWAS summary statistics.

**Supplementary Table 5.** Statistics (Principal Component Analysis) of individual covariate effects on gene expression.

**Supplementary Table 6.** List of genes differentially expressed across brain regions in female or male PTSD vs. controls.

**Supplementary Table 7.** List of genes differentially expressed between males and females at baseline (controls only).

**Supplementary Table 8.** Quantitative PCR validation data in females and males with PTSD.

**Supplementary Table 9.** List of genes differentially expressed across brain regions in MDD vs. control.

**Supplementary Table 10.** List of genes differentially expressed across brain regions in female or male PTSD vs. controls.

